# Sphingomyelin biosynthesis is essential for phagocytic signaling during *Mycobacterium tuberculosis* host cell entry

**DOI:** 10.1101/565226

**Authors:** Patrick Niekamp, Gaelen Guzman, Hans Leier, Ali Rashidfarrokhi, Veronica Richina, Fabian Pott, Joost C. M. Holthuis, Fikadu G. Tafesse

## Abstract

*Mycobacterium tuberculosis* (Mtb) invades alveolar macrophages through phagocytosis to establish infection and cause disease. The molecular mechanisms underlying Mtb entry are still poorly understood. Here, we report that an intact sphingolipid biosynthetic pathway is essential for the uptake of Mtb across multiple phagocytic cell types without affecting other forms of endocytosis. Disrupting sphingolipid production prevents the removal of the inhibitory phosphatase CD45 from the nascent phagosome, a critical step in the progression of phagocytosis. We also show that blocking sphingolipid biosynthesis impairs activation of Rho GTPases, actin dynamics and phosphoinositide turnover at the host-pathogen contact site. Moreover, production of sphingomyelin, not glycosphingolipids, is critical for the uptake of Mtb. Together, our data provide mechanistic insights regarding how Mtb exploits sphingomyelin metabolism to enter host cells, representing a potential avenue for designing host-directed therapeutics against tuberculosis.

## Introduction

*Mycobacterium tuberculosis* (Mtb), the etiologic agent of tuberculosis, remains a significant global health threat. It is estimated that more than two billion people around the world are infected with the latent form of Mtb and about 1.5 million die every year (*1*). The success of Mtb as a pathogen relies on its ability to effectively enter the immune cells and establish its niche by subverting the defense mechanisms of the host (*2, 3*). Elucidating these mechanisms will help us to understand the complex host-pathogen interfaces and design a therapeutic strategy to combat this disease.

Phagocytosis of Mtb by lung resident alveolar macrophages represents the obligate first step in the Mtb infection cycle. Phagocytic cells of the immune system consist predominantly of macrophages, dendritic cells and neutrophils and are responsible for the production of key effector molecules critical for instigating the adaptive immune responses (*4*). Phagocytosis is a complex cellular process that begins with the binding of pathogens to the cell surface through the interaction of pattern recognition receptors (PRRs) with ligands on the surface of the pathogen (*5, 6*). During this process, phagocytes undergo extensive membrane reorganization and cytoskeleton rearrangement at their cell surface to engulf the pathogen. Conceivably, the lipid composition of the plasma membrane plays an essential role in regulating the lateral movement of receptors and signaling molecules to form the phagocytic synapse (*6, 7*). While a great deal of effort has been devoted to understanding the proteins and signaling molecules such as phosphoinositides in this process, the contribution of other membrane lipids such as sphingolipids in Mtb phagocytosis remains unknown.

Sphingolipids are a diverse class of lipids that contain a sphingoid base and are conserved in all eukaryotes. Although their distribution among the various biological organelles is distinct, sphingolipids are mainly enriched at the outer leaflet of the plasma membrane (*8–10*). Therefore, pathogens including Mtb inevitably interact with this class of lipids during phagocytosis. The *de novo* biosynthetic pathway of sphingolipids starts with the condensation of serine and palmitoyl CoA to yield 3-ketodihydrosphingosine, in a reaction catalyzed by serine palmitoyl CoA transferase (SPT) (*9*). Subsequent reactions yield ceramide, the backbone of all sphingolipids, which is then delivered to the Golgi apparatus to be converted to sphingomyelin or glycosphingolipids (Figure 1A) (*9, 10*). These molecules are known mediators of a diverse array of cellular functions and signaling, have critical roles in membrane organization at the cell surface (*11, 12*). Moreover, sphingolipids are essential for the transport of viral glycoproteins from the Golgi apparatus to the cell surface and for the trafficking of bacterial toxins across the endocytic pathway (*13, 14*). Although the type of sphingolipid species and their mechanism of actions remained to be determined, recent studies have shown a role for sphingolipids in *in vivo* clearance of the pathogenic fungi *Candida albicans* through phagocytosis (*15*). Thus far, we know little about whether bioactive and structural sphingolipids play a role in Mtb invasion of phagocytic immune cells.

**Figure 1:**
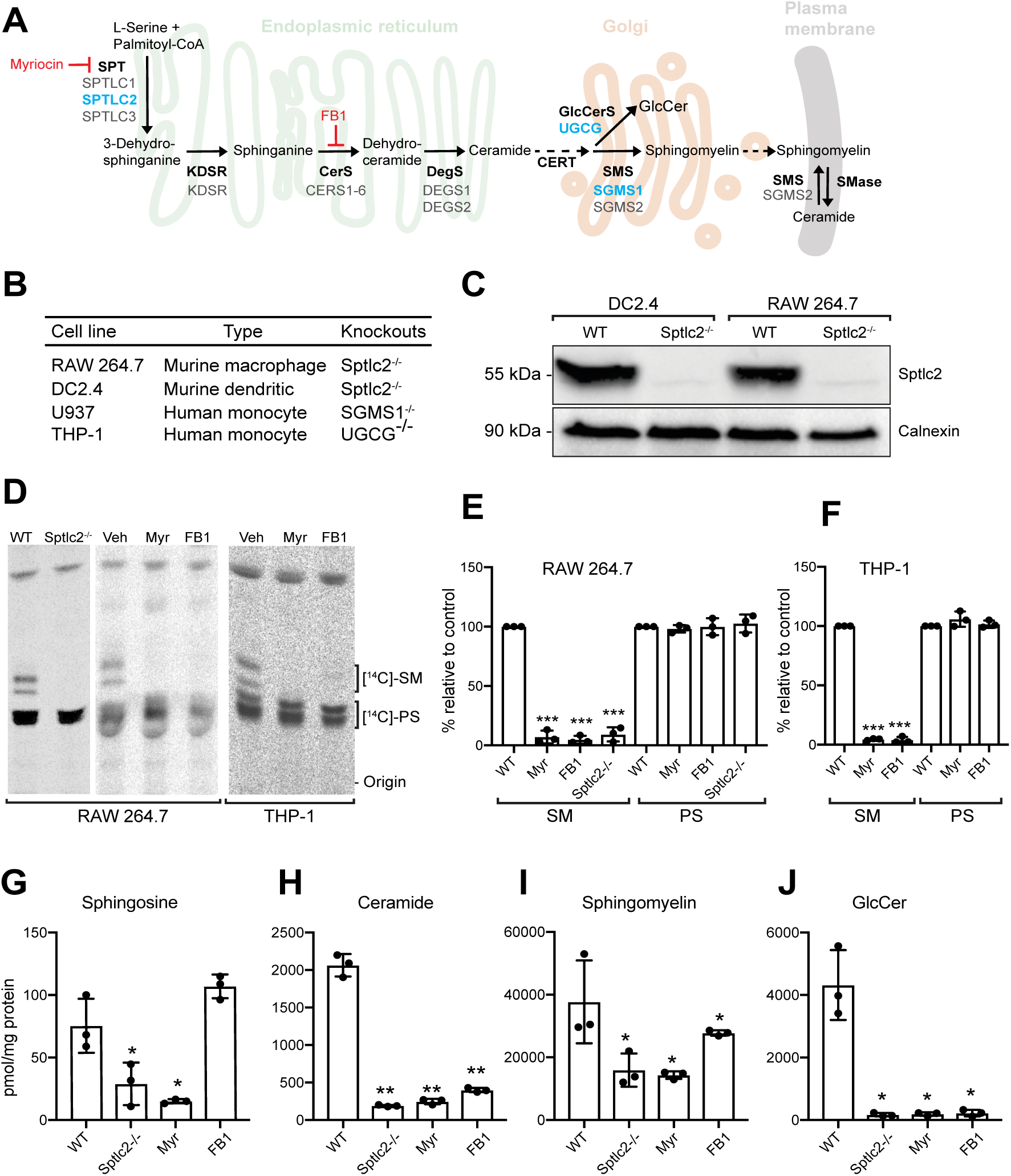
Manipulation of cellular sphingolipid levels using genetics and chemical tools. (A) Simplified overview of sphingolipid biosynthetic pathway. Genes that are deleted using CRISPR/Cas9 system in this study are indicated in blue and the inhibitors used in this study are shown in red. (B) Model phagocyte cell lines used in this study, respective species background, and genetic knockouts available. (C) DC2.4 and RAW 264.7 wild type and Sptlc2^-/-^ cells were processed for immunoblotting with antibodies against Sptlc2 and Calnexin. (D) Control, Sptlc2^-/-^ and inhibitor-treated cells were labeled with the sphingolipid precursor 3-L-[^14^C]-serine, and total lipids were extracted and analyzed by thin layer chromatography (TLC) and autoradiography. (E, F) Quantification of the [^14^C]-SM and [^14^C]-PS signals from [^14^C]-serine labeling of RAW 264.7 and THP-1 cells from TLC shown in *D*. (G-J) Total lipids were extracted from control, Sptlc2^-/-^ and inhibitor-treated cells and lipid profiling was performed using LC/MS. Levels of sphingosine (G), ceramide (H), sphingomyelin (I) and glucosylceramide (GlucCer; J) are shown. All graphs display SD of three independent experiments, and an unpaired t-test was used to analyze the significance of the data. *p<0.05, **p<0.01, *** p < 0.001.

Here we show that Mtb requires host sphingolipids to effectively invade both macrophages and dendritic cells. Perturbation of sphingolipid biosynthesis through CRISPR/Cas9 gene editing, or small molecule inhibitors that target the key enzymes in the sphingolipid biosynthetic pathway, remarkably reduces Mtb uptake in different phagocytic cell types without affecting other forms of receptor-mediated endocytosis. Employing extensive time-lapse live imaging, we determined that sphingolipids are critical in the removal of the inhibitory phosphatase CD45 from the receptor-pathogen contact sites, an essential step in the progression of a phagocytic synapse. We also found that macrophages deficient in sphingolipid biosynthesis are defective in phosphoinositide turnover, F-actin remodeling and activation of the small GTPases Rac1 and CDC42. We further determined that biosynthesis of sphingomyelin, not glycosphingolipids, is critical for the phagocytosis of Mtb by macrophages. These data demonstrate a role for sphingomyelin biosynthesis in an early step of Mtb infection, defining a potential target for host-directed anti-mycobacterial therapeutics.

## Results

### Manipulation of sphingolipid levels in mammalian phagocytes using pharmacological and genetic tools

To define the role of the sphingolipid biosynthetic pathway in phagocytic uptake of Mtb, we sought to inhibit sphingolipid biosynthesis in four different phagocytic cell lines: the murine macrophage cell line RAW 246.7, the murine dendritic cell line DC2.4, and the human monocyte cell lines THP-1 and U937 (Figure 1B). As depicted in Figure 1A, the initial and rate limiting step of sphingolipid synthesis is the production of the sphingoid base through the condensation of serine and palmitoyl-CoA, a reaction performed by the serine palmitoyltransferase (SPT) enzyme complex (*9*). Previously, we reported that genetic ablation of the SPT subunit Sptlc2 blocks the *de novo* synthesis of all sphingolipids, as does treatment with the atypical amino acid myriocin (Myr) (*15*). The fungal toxin fumonisin B1 (FB1) acts several steps downstream in the sphingolipid biosynthetic pathway by blocking the activity of ceramide synthases (*15*). As the central nexus of sphingolipid biosynthesis, ceramide is converted into a variety of complex sphingolipids, including sphingomyelin and glycosphingolipids (*9, 16*).

To verify the reported effects of Myr and FB1 on *de novo* sphingolipid biosynthesis, we monitored the incorporation of 3-L-[^14^C]-serine into sphingolipid species in RAW 246.7, THP-1, DC2.4 and U937 cells after 3 days of drug treatment using thin layer chromatography analysis. Across all four cell lines examined, treatment with Myr or FB1 in each case reduced incorporation of the serine isotope into sphingomyelin by more than 90%; in contrast, incorporation of the isotope into phosphatidylserine was largely unaffected (Figure 1D, 1E and 1F; Supplemental Figure 1). Genetic ablation of Sptlc2 in RAW 246.7 and DC2.4 cells gave the same results (Figure 1C, 1D and 1E; Supplemental Figure 1), in line with our previous findings (*15*).

We next examined the impact of drug treatment and Sptlc2 removal on the steady state sphingolipid levels. Both Myr-treated and Sptlc2^-/-^ cells contained significantly reduced levels of sphingosine, ceramides, sphingomyelin and glucosylceramide in comparison to control cells (Figure 1 G-J). In FB1-treated cells, we observed a reduction in the levels of ceramide, sphingomyelin and glucosylceramide concomitant with an increase in sphingosine levels, as expected (Figure 1G-J) (*17*). The residual sphingolipid levels detected in Sptlc2^-/-^, Myr- and FB1-treated cells are likely due to the activity of sphingolipid salvage pathways, through which cells may acquire sphingolipids from additives in the culture medium – primarily serum. This enables them to bypass the growth defect that is generally observed when sphingolipid biosynthetic mutants are cultured in serum-depleted medium. Indeed, we observed that Sptlc2^-/-^ DC2.4 and Sptlc2^-/-^ RAW 264.7 cells grew equally well in serum-containing medium as their wild type counterparts (Supplementary Figure 2) (*15*).

**Figure 2:**
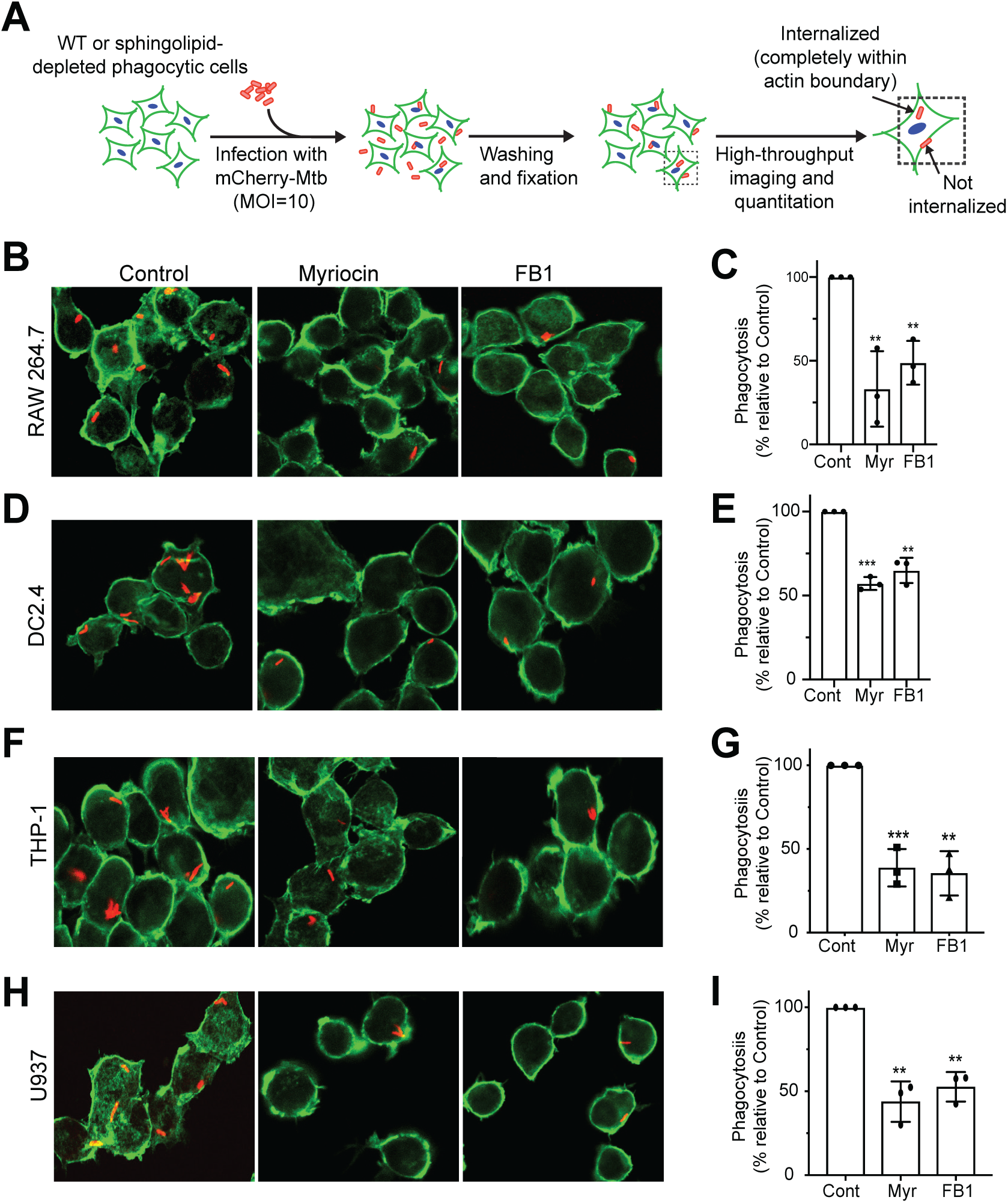
Sphingolipid biosynthesis is required for efficient phagocytosis of Mtb. (A) Overview of the phagocytosis experiment. Cells were infected with an mCherry-expressing Mtb and fixed after 2h. Cell outline was stained with Phalloidin A488 to differentiate between internalized and non-internalized Mtb and the nuclei was stained with DAPI to quantify the number of cells. (B, D) Representative images of Mtb-infected RAW 264.7 (B) and DC2.4 (D) cells. (C, E) Quantification of Mtb uptake in RAW 264.7 (C) and DC2.4 (E) cells. (E, H) Representative images of infected THP-1 (F) and U937 (H) monocyte-derived macrophages. (G, I) Quantification of Mtb uptake in THP-1 (G) and U937 (I) cells. All cell lines were treated for three days with myriocin or FB1. Three independent experiments were performed each in triplicate with at least 1,000 cells quantified per replicate. Mtb uptake was defined using total number of bacteria (mCherry signal) within cellular periphery (Phalloidin-Alexa488 signal) and cellular counts were defined by nuclear stain. Data are mean ± SD. ***p<0.001, two-tailed unpaired t-test. Data are mean ± SD. **p<0.01, ***p<0.001, two-tailed unpaired t-test.

### Cells deficient in sphingolipid biosynthesis display a reduced capacity to phagocytose Mtb

To address whether sphingolipids participate in the phagocytic uptake of Mtb, RAW 264.7, THP-1, DC2.4, and U937 cells were treated with Myr and FB1 for 3 days and then infected with a mCherry-expressing strain of Mtb (Figure 2A). In brief, cells were infected at a multiplicity of infection (MOI) of 10 for 2 h, fixed and then stained with DyLight 488-conjugated phalloidin and DAPI to visualize F-actin and cell nuclei, respectively. Fluorescence microscopy was used to quantify uptake efficiency by automated counting of cell number (by nuclei) and internalized bacteria (red fluorescent particles within green cell boundary). Uptake efficiency is defined as the ratio of the total number of internalized Mtb particles against the total number of identified nuclei. The ratio is reported as a percentage of phagocytosis relative to the control cells. In all four cell models analyzed, we observe that Myr and FB1 treatment caused a significant (40-60%) reduction in Mtb uptake (Figure 2B-I). Similarly, the knockout of Sptlc2 in RAW 264.7 and DC2.4 cells results in a ∼50% reduction in uptake relative to their wild type controls (Figure 3A-D). Collectively, these results indicate that an intact sphingolipid biosynthetic pathway is essential for an efficient uptake of Mtb by professional phagocytes.

**Figure 3:**
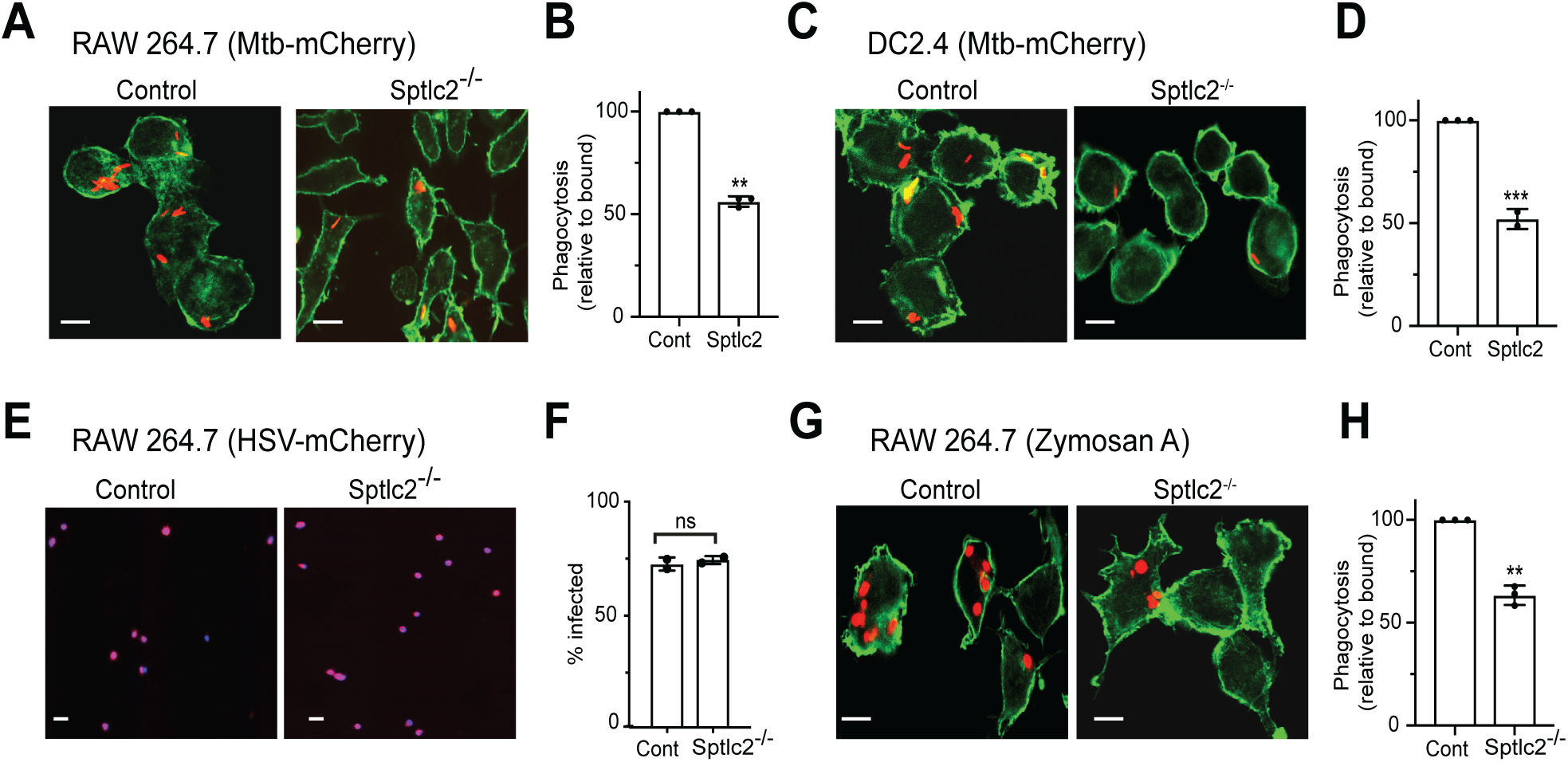
Sphingolipid biosynthesis is required for efficient phagocytosis of Mtb and Zymosan A, but not for HSV entry. (A) Representative images of wild type and Sptlc2^-/-^ RAW 264.7 (A) and DC2.4 (C) cells infected with mCherry-expressing Mtb. (B, D) Quantification of Mtb uptake for experiments performed as in *A* and *C*, respectively. (E) Microscopy images of cells that were infected with HSV-mCherry (MOI of 1). At 12 h post infection, cells were fixed, stained with DAPI and analyzed by microscopy. (F) Quantification of experiments performed as described *E*. (G) Representative images of wild type and Sptlc2^-/-^ RAW 264.7 cells infected with Alexa Fluor 594-conjugated Zymosan A particles. (H) Quantification of Zymosan A uptake performed as in *E*. Three independent experiments were performed each in triplicate with at least 1,000 cells quantified per replicate. Data are mean ± SD. ***p<0.001, two-tailed unpaired t-test. Data are mean ± SD. **p<0.01, ***p<0.001, ns = not significant, two-tailed unpaired t-test.

### Sptlc2^-/-^ cells are permissive to endocytic uptake of Herpes Simplex Virus (HSV)

To determine whether an intact sphingolipid biosynthetic pathway is a general prerequisite for the entry of pathogens in professional phagocytes, we performed parallel infections of Sptlc2^-/-^ and wild type RAW 264.7 cells with a fluorescent reporter strain of Herpes Simplex Virus-1 (HSV-1) encoding a fusion of the viral protein VP26 and mCherry (*18*). HSV-1 is an enveloped virus that gains entry to host cells through the phagocytosis-like process of micropinocytosis; the viral receptor involved is nectin-1, also known as herpesviruses entry mediator (*19*). Previous work revealed that HSV-1 entry is dependent on cholesterol (*20*). At 12 h post-infection, we quantified the rates of HSV-1 infection using high-content imaging as described above. Because the VP26-mCherry reporter construct localizes to the nucleus, we did not stain cells using phalloidin and instead used the co-localization of DAPI with mCherry to determine infection efficiency. However, we observe no significant difference in the HSV-1 infection rates between wild type and Sptlc2^-/-^ cells (Figure 3E and 3F). These data indicate that sphingolipid biosynthesis is dispensable for HSV-1 entry and support a more specific role of sphingolipids in the phagocytic uptake of Mtb.

### Sptlc2^-/-^ cells fail to exclude CD45 from the contact site of pathogen-mimicking particles

We next investigated whether blocking sphingolipid biosynthesis also affects the phagocytic uptake of Zymosan A particles. These pathogen-mimicking particles have been shown to engage the C-type Lectin receptor Dectin-1 on professional phagocytes (*15, 21*). Dectin-1 is one of several pathogen recognition receptors that engage Mtb and initiate phagocytic signaling to facilitate host cell entry (*22*). Interestingly, Sptlc2^-/-^ RAW 264.7 cells displayed a ∼40% reduction in the phagocytic uptake of Zymosan A relative to wild type cells, analogous to Mtb uptake (Figure 3G and 3H). This led us to use Zymosan A as a model particle in subsequent experiments, in which we aimed to define which step in phagocytosis relies on an intact sphingolipid biosynthetic pathway.

The engagement of pathogen recognition receptors with their ligands initiates receptor clustering at the phagocytic cup, followed by a lateral segregation of the inhibitory phosphatase CD45 (*7*). Because sphingolipids have been implicated as active participants in the organization of signaling complexes at the cell surface (*23*), we reasoned that sphingolipid levels at the plasma membrane may be critical for the aggregation of Dectin-1 and/or exclusion of CD45 from that pathogen contact site. To experimentally address this idea, we transfected wild type and Sptlc2^-/-^ RAW 264.7 cells with a GFP-tagged Dectin-1 construct and assessed the co-localization of the fluorescent receptor with CD45 following treatment with non-fluorescent Zymosan A particles. In both wild type and Sptlc2^-/-^ cells, Dectin-1 aggregated at the Zymosan A contact site within two minutes after particle exposure, indicating that sphingolipid biosynthesis is dispensable for ligand-induced receptor clustering. As expected, in wild type cells the inhibitory phosphatase CD45 was efficiently segregated from the Dectin-1 clusters at the particle contact site (Figure 4A upper panels and B) (*7*). In contrast, CD45 failed to segregate from the Dectin-1 clusters in Zymosan A-treated Sptlc2^-/-^ cells (Figure 4A lower panels and B).

**Figure 4:**
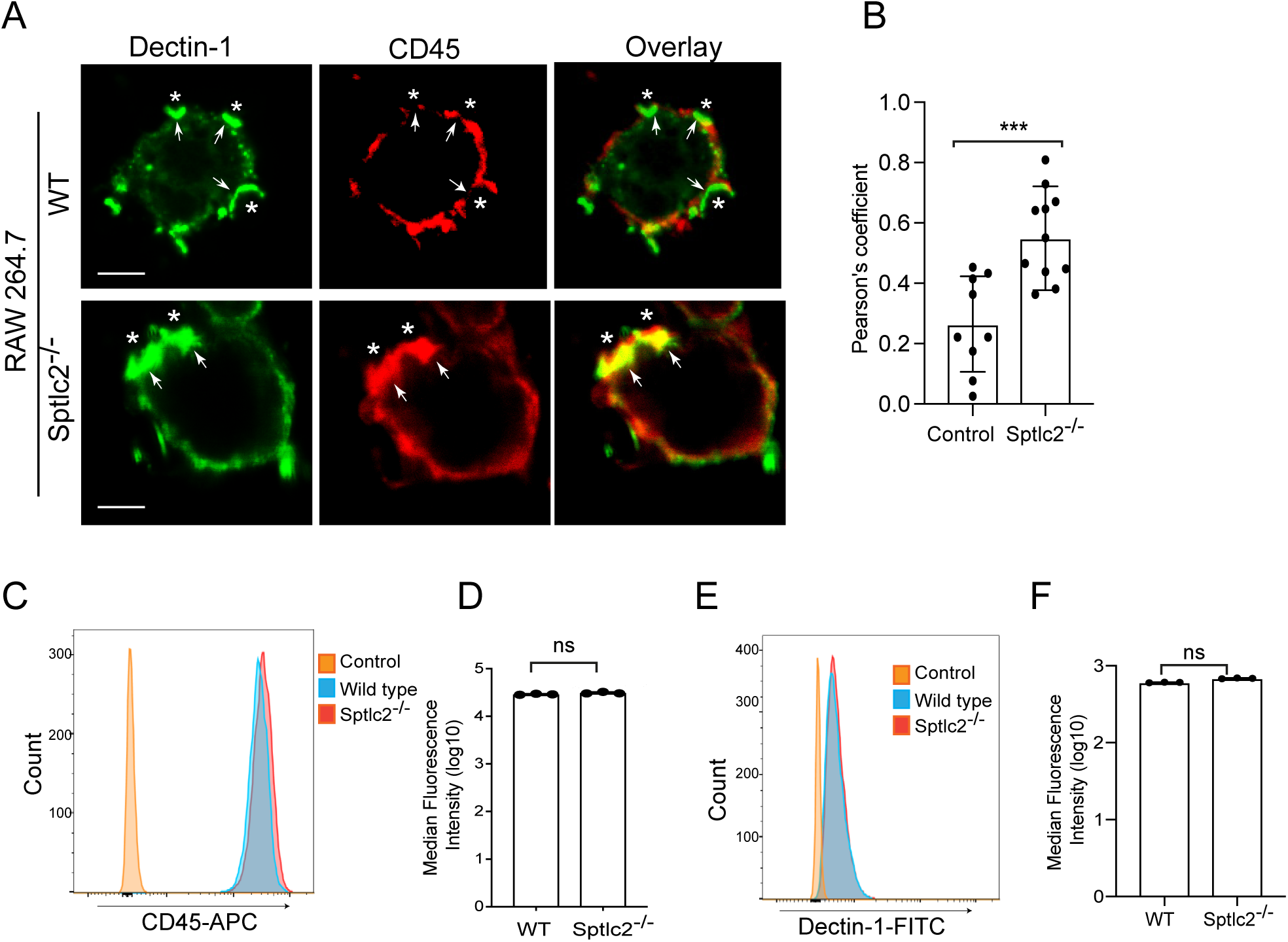
Exclusion of CD45 by Dectin-1 from the cup formation site is impaired in Sptlc2^-/-^ cells. (A) Confocal microscopy of GFP-tagged Dectin-1-expressing RAW 264.7 macrophages stimulated with Zymosan A for 2 min and stained for CD45 (red). Zymosan particles are shown as star (*) at the phagocytic cup. (B) Colocalization measurement of CD45 and Dectin-1 at the site of cup formation as presented by Pearson’s colocalization coefficient. (C, E) Flow cytometry of wildtype and Sptlc2^-/-^ RAW 264.7 cells that were stained for CD45 and Dectin-1at the cell surface. (D, F) Graphs showing quantification of cell surface expression of the receptors described in *C* and *E*, respectively. All data are representative of 3 independent experiments performed in triplicate. Data are mean ± SD. ns = not significant, two-tailed unpaired t-test.

To address whether a block in sphingolipid biosynthesis affects the overall surface expression of CD45 and Dectin-1, wild type and Sptlc2^-/-^ RAW 264.7 cells were stained with APC-conjugated anti-CD45 and FITC-conjugated anti-Dectin-1 antibodies and subjected to analytical flow cytometry. We found no differences in the overall fluorescent signals for both markers between wild type and Sptlc2^-/-^ cells (Figure 4C-F), indicating that the cell surface levels of endogenous Dectin-1 and CD45 are unaffected by blocking sphingolipid biosynthesis. From this we conclude that an intact sphingolipid biosynthetic pathway is required to enable a proper displacement of CD45 from the site of Dectin-1 clustering, a step critical for the initiation of phagocytic synapse formation.

### Activation of Rho GTPases Rac1 and Cdc42 is impaired in Sptlc2^-/-^ cells

Phagocytosis is strictly regulated through a signaling network that involves the recruitment of pathogen receptors and Rho GTPases to the particle contact site. Activated Rho GTPases promote the conversion of phosphatidylinositol-4-phosphate (PI4P) to phosphatidylinositol-4,5-*bis*phosphate (PIP2) by stimulating phosphatidylinositol 5-kinase (PI5K). The local accumulation of PIP2 is essential to initiate the polymerization of filamentous actin (F-actin), a process that drives extension of the leading edges of the nascent phagosome around the particle (*24*). However, to complete the engulfment and internalization of the particle, F-actin must disassemble at the base of the phagocytic cup. This is achieved through a local conversion of PIP2 into phosphatidylinositol-3,4,5-*tris*phosphate (PIP3) by the phosphatidylinositol 3-kinase (PI3K; Figure 5A) (*4, 24*).

**Figure 5.**
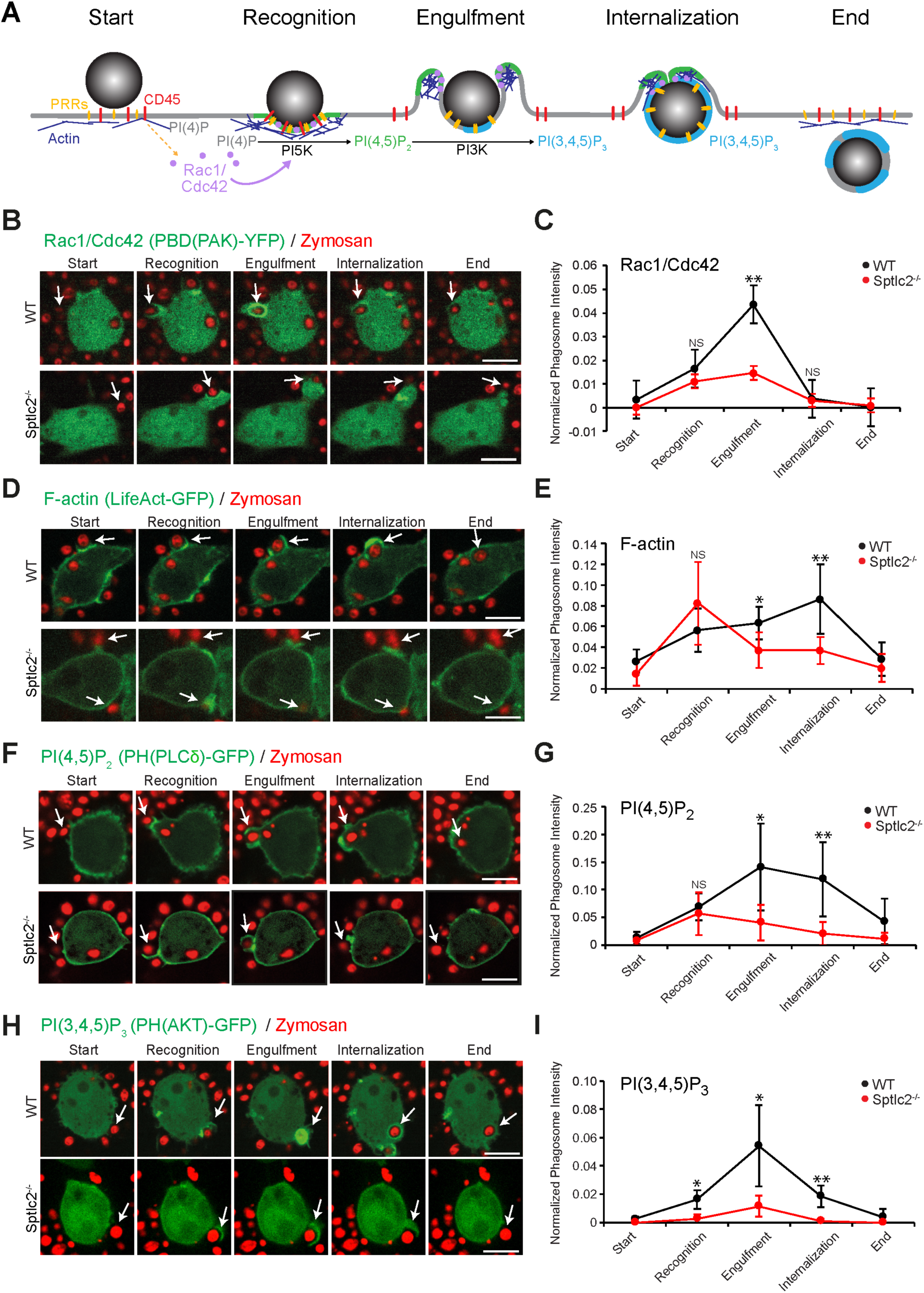
Sphingolipids are essential for efficient phosphoinositide turnover at the phagocytic cup. (A) Schematic representation of the phagocytosis process. Upon pathogen engagement, pattern recognition receptors (PRRs) cluster and displace CD45 phosphatase, thereby initiating the activation of the Rho GTPases Cdc42 and Rac1, which in turn activate PI5K. This leads to a spatial production of PIP2 and F-actin assembly. To permit phagosome progression, PI3K orchestrates the turnover of PIP2 to PIP3, leading to F-actin disassembly at the base of the phagosome. (B) Time-lapse confocal images of wild type and Sptlc2^-/-^ RAW 264.7 cells transfected with PAK(PBD)– GFP and infected with Zymosan Alexa Flour 594-conjugate at a MOI of 10. (C) Quantification of the PAK(PBD) fluorescence intensity in the course of the formation of a phagocytic cup (shown in arrow) as performed in *B*. (D) Time-lapse confocal images of wild type and Sptlc2^-/-^ RAW 264.7 cells transfected with LifeAct-GFP and infected with Zymosan Alexa Flour 594-conjugate at a MOI of 10. (E) Quantification of the LifeAct fluorescence intensity in the course of the formation of a phagocytic cup (shown in arrow) as performed in *D*. (F) Time-lapse confocal images of wild type and Sptlc2^-/-^ RAW 264.7 cells transfected with PH(PLCδ)-GFP and infected with Zymosan Alexa Flour 594-conjugate at a MOI of 10. (G) Quantification of the PH(PLCδ) fluorescence intensity in the course of the formation of a phagocytic cup (shown in arrow) as performed in *F*. (H) Time-lapse confocal images of wild type and Sptlc2^-/-^ RAW 264.7 cells transfected with PH(AKT)-GFP and infected with Zymosan Alexa Flour 594-conjugate at a MOI of 10. (I) Quantification of the PH(AKT) fluorescence intensity in the course of the formation of a phagocytic cup (shown in arrow) as performed in *H*. Accumulation of GFP signal to site of particle binding was quantified by the ratio of increased GPF signal intensity at the phagosome versus the GFP signal across entire cell at the indicted points. Intensity profiles of 5-10 cells are represented. Data are mean ± SD. *p<0.05, **p<0.01, two-tailed unpaired t-test, ns = not significant.

To determine whether blocking sphingolipid biosynthesis perturbs phagocytic signaling events downstream of pathogen receptor activation, we first focused on the activation and localization of the Rho GTPases Rac1 and Cdc42 in Zymosan A-treated wild type and Sptlc2^-/-^ RAW 264.7 cells. Visualization of activated Rho GTPases was enabled by transfecting cells with a fluorescent biosensor comprising YFP fused to the p21-binding domain of the p21-activated kinase PAK (PBD(PAK)-YFP) (*24, 25*). Twenty-four hours post transfection, cells were infected with Zymosan A particles at an MOI of 10 and subjected to live cell imaging. In wild type cells, we observed a sharp increase in PBD(PAK)-YFP fluorescence at the particle contact site. Fluorescence levels peaked during particle engulfment, corresponding to the expected pattern of pathogen-induced Rac1/Cdc42 activation (Figure 5B and 5C; Movie 1). In Sptlc2^-/-^ cells, we also observed an increase in PBD(PAK)-YFP fluorescence upon particle recognition. However, the PBD(PAK)-YFP fluorescence intensity failed to reach the levels detected in Zymosan A-treated wild type cells and faded out prior to a successful particle internalization, suggesting that Sptlc2^-/-^ cells are impaired in particle-induced activation of Rho GTPases (Figure 5B and 5C; Movie 2).

We next visualized F-actin dynamics in Zymosan A-treated wild type and Sptlc2^-/-^ cells using LifeAct-GFP (*26*). In wild type cells, LifeAct-GFP was readily recruited to the particle contact site. During particle engulfment, LifeAct-GFP accumulated at the leading edges of the phagosome while disappearing from the base of the phagocytic cup (Figure 5D and 5E; Movie 3). In Sptlc2^-/-^ cells, LifeAct-GFP was readily recruited to the particle contact site yet remained associated with the base of the phagocytic cup and failed to accumulate at the leading edges of the nascent phagosome (Figure 5D and 5E; Movie 4). From this we conclude that a block in sphingolipid biosynthesis severely disrupts phagocytosis-associated F-actin dynamics, presumably causing the uptake defect.

### Phosphoinositide turnover is impaired in Sptlc2^-/-^ cells

As PIP2 and PIP3 are key regulators of F-actin dynamics during phagocytosis (Figure 5A), we next monitored the spatial distribution of these phosphoinositides in Zymosan A-treated wild type and Sptlc2^-/-^ cells. To this end, cells were first transfected with a GFP-fusion of the plekstrin homology (PH) domain of the phospholipase C-delta (PH(PLCδ)-GFP), which serves as a fluorescent biosensor for PIP2 (*4, 27*). In Zymosan A-treated wild type cells, PH(PLCδ)-GFP was readily concentrated at the particle contact site and preferentially accumulated at the leading edges of the nascent phagosome during particle engulfment (Figure 5F and 5G; Movie 5). In Sptlc2^-/-^ cells, however, PH(PLCδ)-GFP was readily recruited to the contact site but remained associated with the base of the phagocytic cup, analogous to what we observed with the LifeAct-GFP probe. This suggests PIP2 synthesis is initiated upon Zymosan A binding, but that PIP2 turnover at the base of the phagocytic synapse is impaired in Sptlc2^-/-^ cells (Figure 5F and 5G; Movie 6).

To check whether a block in sphingolipid biosynthesis affects the turn-over of PIP2 into PIP3 in Zymosan-A-treated cells, wild type and Sptlc2^-/-^ RAW 264.7 cells were transfected with a GFP-fusion of the PH domain of Akt (PH(AKT)-GFP), which serves as a fluorescent biosensor for PIP3 (*4, 27*). In wild type cells, PH(AKT)-GFP was readily recruited to the particle contact site, and its membrane association increased sharply during particle engulfment (Figure 5H and 5I; Movie 7). In contrast, Sptlc2^-/-^ cells almost completely failed to mobilize PH(AKT)-GFP at the particle contact site (Figure 5F and 5G; Movie 8). Together, these results indicate that blocking sphingolipid biosynthesis disrupts phosphoinositide metabolism at phagocytic synapses in Zymosan A-treated macrophages, notably at the level of PIP2-to-PIP3 conversion, a step known to be critical for the internalization of pathogenic bacteria by professional phagocytes (*24*).

### Production of sphingomyelin, not glycosphingolipids, is critical for efficient Mtb uptake

Both sphingomyelin and glycosphingolipids are highly enriched in the exoplasmic leaflet of the plasma membrane of mammalian cells. Therefore, we next addressed whether efficient phagocytic uptake of Mtb relies on the production of sphingomyelin, glycosphingolipids, or both. Bulk production of sphingomyelin is mediated by the enzyme sphingomyelin synthase 1 (SGMS1) in the *trans*-Golgi whereas glycosphingolipid production is strictly dependent on the enzyme UDP-glucose ceramide glycosyltransferase (UGCG) in the *cis*-Golgi (*9, 28*). We used CRISPR/Cas9 gene editing to ablate UGCG expression in THP-1 cells (Supplemental Figure 3), and the lab of Dr. Virriera-Winter kindly provided us with SGMS1 knockout U937 cells (*14*). Strikingly, SGMS1 knockout resulted in a major (∼60%) reduction in the phagocytic uptake of mCherry-expressing Mbt by U937 cells (Figure 6A and 6B), similar to what we observed in myriocin or FB1-treated U937 cells (Figure 2H and 2I). In contrast, genetic ablation of UGCG had no impact on Mtb uptake by THP-1 cells (Figure 6C and 6D), even though phagocytosis of Mtb in THP-1 cells can be readily inhibited by myriocin and FB1 (Figure 2F and 2G). Together, these data indicate that an efficient uptake of Mtb by professional phagocytes is critically dependent on an intact sphingomyelin biosynthetic pathway whereas the production of glycosphingolipids is dispensable for this process.

**Figure 6:**
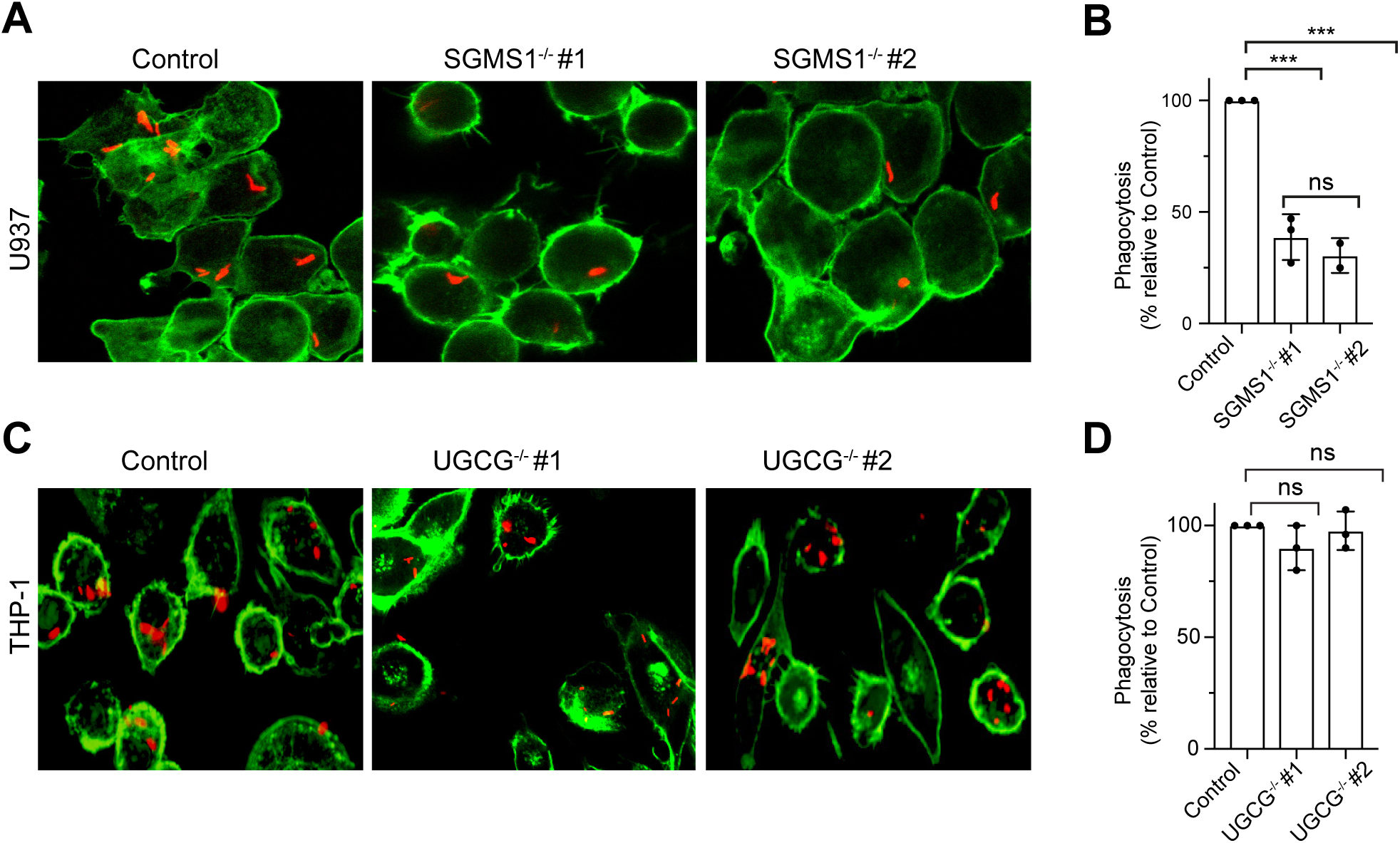
Production of sphingomyelin, not glycosphingolipids, is critical for efficient Mtb uptake. (A) Representative images of U937 wild type and two different SGMS1 knockout cell lines infected with mCherry-expressing Mtb for 2h. (B) Quantification of Mtb uptake for experiments described as in *A*. (C) Representative images of THP-1 wild type and two different UGCG knockout cell lines infected with mCherry-expressing Mtb for 2h. (D) Quantification of Mtb uptake for experiments described as in *C*. U937 and THP-1 cells were activated with 50nM PMA for 24 h before Mtb infection. Three independent experiments were performed each in triplicates with at least 1,000 cells quantified per replicate. Data are mean ± SD. ***p<0.001, ns = not significant, two-tailed unpaired t-test.

## Discussion

In this study, we report that the invasion of phagocytes by Mtb is critically dependent on an intact sphingolipid biosynthetic pathway. Blocking sphingolipid production does not affect macropinocytosis of Herpes Simplex Virus, indicating that phagocytes remain susceptible to infection by pathogens that utilize other routes of cellular uptake. Strikingly, sphingolipid biosynthetic Sptlc2^-/-^ mutant cells display normal pathogen-induced clustering of phagocytic receptors, yet fail to exclude the regulatory phosphatase CD45 from the phagocytic synapse. We also show that key steps in the downstream signaling cascade, notably Rho GTPase activation and phosphoinositide turnover, are significantly impaired in the mutant cells, leading to perturbations in actin dynamics and defective phagosome formation. Blocking production of sphingomyelin, not glycosphingolipids, suffices to disrupt phagocytic uptake of Mtb. While it remains to be determined if this class of lipids plays a role in intracellular bacterial replication and disease outcome *in vivo*, our findings disclose a vital role of sphingomyelin metabolism in an early step of Mtb invasion.

A major finding of this work is that macrophages defective in sphingolipid biosynthesis are unable to segregate CD45 from pathogen-induced clusters of Dectin-1 at the phagocytic cup. Dectin-1 belongs to a family of phagocytic cell surface receptors that initiate internalization of pathogens, including Mtb (*7, 29*). Upon pathogen engagement, these receptors cluster in the lateral plane of the membrane, bringing their cytosolic domains in close apposition (*7*). The cytosolic domain of Dectin-1 contains an immunoreceptor tyrosine-based activation motif (ITAM), which becomes phosphorylated by Src and Syk family kinases upon receptor coalescence (*30*). Importantly, segregation of the regulatory tyrosine phosphatase CD45 from the site of Dectin-1 clustering is a crucial step in the amplification of a downstream phosphorylation cascade (*7*), which eventually culminates in the recruitment of phosphoinositide-modifying enzymes including the phosphoinositide 3-kinase (PI3K) (*4, 31*). Thus, our finding that Sptlc2^-/-^ cells fail to generate PIP3 at the phagocytic cup might be due to an impaired phagocyte receptor signaling caused by a defective segregation of CD45. PI3K activation and formation of PIP3 at the pathogen engagement site are necessary for late stages of pseudopodia progression, the clearance of actin at the base of the phagocytic cup, and phagosome closure (*4*). Collectively, our data suggest that blocking sphingolipid biosynthesis affects the phagocytic uptake of Mtb by precluding a proper activation of phagocyte receptors during pathogen engagement.

Another key observation is that production of sphingomyelin rather than sphingolipid biosynthesis *per se* is critical for efficient phagocytic uptake of Mtb. Sphingomyelin is a dominant sphingolipid in the plasma membrane of mammalian cells. Owing to its saturated nature and hydrogen-bonding capacity, sphingomyelin enables the formation of cholesterol/sphingolipid-enriched microdomains (*32*). In this capacity, sphingomyelin may serve a critical role in the lateral organization and activation of signaling complexes that promote Mtb invasion, consistent with prior reports that cholesterol is essential for Mtb entry into macrophages (*33*). However, an alternative scenario is that a sphingomyelin-derived metabolite, ceramide, facilitates the phagocytic uptake of Mtb. Previous work revealed that hydrolysis of cell surface sphingomyelin by acid sphingomyelinase mediates the invasion of gram-negative pathogens like *Neisseria gonorrhoeae* and *Pseudomonas aeruginosa* in mammalian cells (*34, 35*). Activation of acid sphingomyelinase at the pathogen contact site appears to trigger local assembly of ceramide-enriched signaling platforms that initiate internalization (*36*). While sphingomyelinases and ceramides have also been implicated in mycobacterial infection (*37, 38*), the underlying molecular principles remain to be explored.

In sum, we show that Mtb relies on an intact sphingomyelin biosynthetic pathway to efficiently invade professional phagocytes. While the latter process is critical for the clearance of pathogens, Mtb harnesses phagocytosis to gain entry to the host cells and establish long-term survival. By providing fresh insights into the underlying mechanisms, our study makes a significant contribution to the development of new strategies to fight this ancient pathogen.

## Materials and Methods

### Reagents

Myriocin and FB1 were purchased from Cayman Chemical Company. 3-L-[^14^C]-serine was purchased from American Radioactivity. Phorbol 12-myristate 13-acetate (PMA) was purchased from BioLegend. 4% paraformaldehyde was purchased from Fisher Scientific. DyLight® 488 Phalloidin was purchased from Cell Signaling Technology, and Alexa Fluor 647-conjugated Zymosan A (*S. cerevisiae*) BioParticles were purchased from Thermo Fisher. Mouse Monoclonal antibody against GAPDH (ab8245) was purchased from Abcam. Anti-rabbit IgG HRP-linked antibody and anti-mouse IgG HRP-linked antibody were purchased from Cell Signaling. APC-conjugated, rat monoclonal antibody against CD45 antibody (Clone 30-F11) was purchased from Biolegend. FITC-conjugated, rat monoclonal antibody against Dectin-1 (Clone 2A11) was purchased from AbD Serotec. Zombie Aqua™ Fixable Viability Kit (part #77143) was purchased from BioLegend.

### Plasmids

The plasmids used for generation of lentivirus consists of the pCDH-CMV(-)-sgRNA plasmid, the psPAX2 (packaging) and pMD2.G (envelope) plasmids. pCW-Cas9 lentivirus was purchased from Addgene. The pMXsIP-Dectin-1-GFP was kindly provided by the lab of Dr. Hidde Ploegh (Boston Children’s Hospital, Boston, MA) (*7*). Active (GTP-bound) Rac1/Cdc42 was detected with PAK(PBD)-YFP, a plasmid encoding the PBD of PAK fused to YFP was obtained from Addgene (*25*). PI(4,5)P2 was detected with PH(PLCδ)-GFP, a plasmid encoding the PH domain of PLCδ fused to GFP. PI(3,4,5)P3 levels were monitored by PH(AKT)-GFP, a plasmid encoding the PH domain of AKT fused to GFP (*39*). These two fusion constructs were kindly provided by the lab of Dr. Hidde Ploegh (Boston Children’s Hospital, Boston, MA).

### Cell culture and small molecule inhibitor treatment

THP-1, U937, and DC2.4 cells were routinely cultured in RPMI medium supplemented with 10% FBS (Seradigm) and 1% Pen Strep (Gibco). RAW 264.7 cells were cultured in DMEM supplemented with 10% FBS and 1% PenStrep. All cell lines were cultured at 37 °C and 5% CO_2_. In plating adherent cells (RAW 264.7 and DC2.4), cells were lifted using 0.25% trypsin solution (Gibco) and counted using hemocytometer, with Trypan Blue solution (Gibco) for counterstain. THP-1 and U937 cells were differentiated to macrophages by 50ng/ml PMA for 1 day in RPMI medium supplemented with 10% FBS, after which the cells were treated with 5µM myriocin or FB1 for 3 days in RPMI supplemented with 10% FBS and 1%Pen Strep. RAW 264.7 and DC2.4 cells were treated similarly with myriocin and FB1 – however, 24-hours before infection, cells were lifted and counted, and seeded at appropriate density in 96-well plates or on coverslips. Biological replicates represent independent treatments with chemical inhibitors and infections on separate days.

### Metabolic labeling and thin layer chromatography

5×10^5^ cells were seeded into a 6-well plate. Cells were labeled with 1µCi/ml of 3-L-[^14^C]-serine for 4 hours in Opti-MEM supplemented with the appropriate inhibitors. Cells were then washed two times with PBS and lipid extraction was done following the Bligh and Dyer method (*40*). The methanol/chloroform-lipid extracts were dried by nitrogen gas. Dried lipids were re-dissolved in several drops of chloroform/methanol (1:2, vol/vol) and loaded on a TLC plate. Lipids were separated by developing the TLC plate first in acetone and then in a mixture of chloroform, methanol and 25% ammonia solution (50:25:6, vol/vol/vol). Radiolabeled lipids were detected on a Storm 825 Phosphor-Imager (GE Healthcare).

### Lipodomics

LC-MS/MS analysis was performed as described previously (*41*). Lipids were extracted from lysed RAW 246.7 cells according to 50µg of protein by chloroform/methanol extraction1. Before the extraction a standard mix containing Sphingosine (d17:1), Ceramide (d18:1/17), Glycosyl(ß) (C12 Cer) and Sphingomyelin (17:0) was spiked into each sample for normalization and quantification. The dried lipid films were dissolved in mixture of mobile phase A (60:40 water/acetonitrile, including 10 mM ammonium formate and 0.1% formic acid) and mobile phase B (88:10:2 2-propanol/acetonitrile/H20, including 2 mM ammonium formate and 0.02% formic acid) with a ratio of 65:35. HPLC analysis was performed using a C30 reverse-phase column. (Thermo Acclaim C30, 2.1 × 250 mm, 3 µm, operated at 50° C; Thermo Fisher Scientific) connected to an HP 1100 series HPLC (Agilent) HPLC system and a QExactivePLUS orbitrap mass spectrometer (Thermo Fisher Scientific) equipped with a heated electrospray ionization (HESI) probe. The elution was performed with a gradient of 45 min; during 0–3 min, elution starts with 40% B and increases to 100%; in a linear gradient over 23 mins. 100% B is maintained for 3 mins. Afterwards solvent B was decreased to 40% and maintained for another 15 min for column re-equilibration. The flow-rate was set to 0.1 ml/min. MS spectra of lipids were acquired in full-scan/data-dependent MS2 mode. The maximum injection time for full scans was 100 ms, with a target value of 3,000,000 at a resolution of 70,000 at m/z 200 and a mass range of 200–2000 m/z in both, positive and negative mode. The 10 most intense ions from the survey scan were selected and fragmented with HCD with a normalized collision energy of 30. Target values for MS/MS were set at 100,000 with a maximum injection time of 50 ms at a resolution of 17,500 at m/z 200. To avoid repetitive sequencing, the dynamic exclusion of sequenced lipids was set at 10 s. Peaks were analyzed using the Lipid Search algorithm (MKI, Tokyo, Japan). Peaks were defined through raw files, product ion and precursor ion accurate masses. Candidate molecular species were identified by database (>1,000,000 entries) search of positive (+H+; +NH4+) ion adducts. Mass tolerance was set to five ppm for the precursor mass. Samples were aligned within a time window and results combined in a single report. From the intensities of lipid standards and lipid classes absolute values for each lipid in pmol/mg protein were calculated.

### Generation of lentiviruses

Lentivirus production was performed as described previously (*42*). Briefly, lentiviruses were produced by co-transfection of HEK 293T cells with the lentiviral vectors that contain our gene of interest (pCDH-sgRNA or pCW-Cas9) and the packaging plasmids (psPAX2 and pMD2.G). Transfection was performed with Lipofectamine 3000 (Thermo Fisher) according to manufacturer’s instructions. Cells were cultured in DMEM supplemented with 10%FBS and the growth medium was replaced after 6 hours. 48 hours after transfection, lentivirus-containing supernatants were harvested, centrifuge for 5min at 1250 rpm and filtered through a 0.45µm filter.

### Generation of CRISPR/Cas9-mediated knockout cell lines

CRISPR/Cas9-mediated genome-editing for genes involved in the sphingolipid pathway were performed as described previously (*15*). Briefly, to generate Sptlc2^-/-^ knockout RAW 264.7 cells were infected with lentivirus (pCW-Cas9) encoding Cas9 cDNA, and were cultured in media containing 2 µg/mL of puromycin (Sigma Aldrich). The following two seed sequences (CRISPR target sequences) preceding PAM motifs found in the open reading frame of Sptlc2 gene were used: Sptlc2 #1 GAACGGCTGCGTCAAGAAC; Sptlc2 #2: AGCAGCACCGCCACCGTCG. The CRISPR target sequence for UGCG was as follows: GCTGTGGCTGATGCATTTCA. Potential off-target effects of the seed sequence were evaluated using the NCBI Mus musculus Nucleotide BLAST. Generation of CRISPR/Cas9-mediated Sptlc2-knockout RAW 264.7 cell line and UGCG-knockout THP-1 cell line were performed as previously described (*42*). Briefly, CRISPR gBlock was designed to clone into the restriction enzymatic site NheI/BamHI of pCDH- CMV(-) (SBI; CD515B-1) as follows:

cacagtcagacagtgactcaGTGTCACAgctagcTTTCCCATGATTCCTTCATATTTGCATATACGATACAAGGCTGTTAGAGAGATAATTAGAATTAATTTGACTGTAAACACAAAGATATTAGTACAAAATACGTGACGTAGAAAGTAATAATTTCTTGGGTAGTTTGCAGTTTTAAAATTATGTTTTAAAATGGACTATCATATGCTTACCGTAACTTGAAAGTATTTCGATTTCTTGGCTTTATATATCTTGTGGAAAGGACGAAACACCGnnnnnnnnnnnnnnnnnnnGTTTTAGAGCTAGAAATAGCAAGTTAAAATAAGGCTAGTCCGTTATCAACTTGAAAAAGTGGC ACCGAGTCGGTGCTTTTTTTggatccTGTGCACAgtcagtcacagtcagtctac (n: CRISPR target sequences).

The gBlock was then digested using the restriction enzymes NheI and BamHI and ligated into pCDH-CMV(-) vector that was linearized by digesting with the same restriction enzyme. The Cas9-inducible cells were infected with lentivirus carrying pCDH-CMV(-)-sgRNA, and were cultured in media containing 250 µg/mL of hygromycin B (Life Technology). To induce expression of Cas9, cells were treated with 1 µg/mL of doxycycline (Clontech) for 3–5 days.

Clonal selection was performed by single cell dilution on 96-well plates. The individual colonies were collected and the expression of Sptlc2 was examined by western-blotting using Sptlc2 antibody. U937 SGSM1^-/-^ cells were kindly provided by Dr. Arturo Zychlinsky (Max Planck Institute of Biochemistry, Berlin, Germany) and the CRISPR/Cas9-mediated knock out cell lines were established as described in Winter et al., 2016 (*14*).

### Culturing of Mtb

Culturing and infection of Mtb was conducted in a Biosafety-level 3 laboratory following general safety guidelines. Mtb strain H37R, expressing constitutively mCherry were obtained from the Fortune lab (Ragon Institute, Cambridge, MA). It was cultured at 37°C in 7H9 Broth medium supplemented with 50µg/ml hygromycin B. The density of the bacteria at the time of infection is between OD600 of 0.6 - 0.8.

### Mtb uptake assay

For infection of THP-1 and U937 cells, the cells were seeded into a SensoPlate™ 96-Well Glass-Bottom Plates (Greiner Bio-One) at a density of 1.5×10^4^ cells per well in RPMI supplemented with 10% FBS, 1% PenStrep and 50ng/ml PMA at 37°C. One day after differentiation, cells were treated for 3 days with 5µM myriocin or 15µM FB1. These cells were then infected with a multiplicity of infection (MOI) of 10 in RPMI supplemented with 10% FBS for 2 hours at 37°C. For infection of RAW 264.7 and DC2.4 cells, 2.0×10^4^ cells per well were seeded into a SensoPlate™ 96-Well Glass-Bottom Plates (Greiner Bio-One). 24 hour later, the cells were infected with an MOI of 10 in RPMI supplemented with 10% FBS for 2 hours at 37°C. Cells were washed with PBS, and then fixed with 4% PFA over night at 4°C.

### Phagocytosis measurement assay

Fixed cells were washed twice with PBS and incubated for 15 min in Permeabilization Buffer (0.1% Triton X-100 and 1%BSA). Cells were stained with Phalloidin-Alexa 488 (Thermo Fisher) at a final concentration of 33nM in permeabilization buffer for 60 min and washed one time with PBS. After the Phalloidin-staining, cells were stained with DAPI for 10 mins and washed 3 times with PBS. Cells were then imaged using the Keyence BZ-X700 with a PlanFluor 20x objective. The parameters for imaging were kept the same for each sample. For image analysis the Keyence BZ-X Analyzer software was used. In brief: The outline of the cells was determined via the Alexa488 signal (Phalloidin staining) and the number and area of mCherry (Mtb) signal in the Alexa488 signal was detected by the program. In a second round the number of cells was determined by counting the nuclei via the DAPI staining. Number of bacteria (mCherry-signal) inside the cell was divided by the number of cells (DAPI signal) to identify the phagocytosis efficiency.

### Herpes Simplex Virus-1 infection

HSV-1-mCherry reporter virus was kindly provided by the lab of Dr. David Johnson (Oregon Health and Science University, Portland, Oregon). Twenty-four hours prior to infection, 1×10^4^ wild type and Sptlc2 knockout RAW 264.7 cells were seeded on glass coverslips in a 12-well plate. Cells were equally infected at an MOI ∼1, and incubated for 12 hours at 37°C and 5% CO_2_. Coverslips were fixed for 20 mins in 4% paraformaldehyde, stained with DAPI and then imaged on a Keyence BZ-X700. Infection rates were calculated by dividing the mCherry expressing cells with the total number of cells (DAPI signal).

### Live imaging of phagocytosis

For transient transfection, RAW 264.7 cells were seeded into a SensoPlate™ 96-Well glass-bottom plates at a cell number of 1.0×10^4^ cells/well 24 hours prior transfection. Transfection was conducted with Lipofectamine 3000 according to manufacturer’s instructions. For imaging, the growth medium was aspirated and replaced with RPMI without phenol red containing Zymosan A Bioparticles Alexa595 at an MOI of 10. The 96-well plates were centrifuged at 1000rpm for one minute. The cells were imaged with the SDC microscope (Nikon) and 3-4 different spots per well were imaged at the same time for 60 min with capture intervals of 15 sec. The parameters for imaging were kept the same for each sample.

### Live image analysis (quantification)

Of live cell images, individual images were selected for further analysis if they captured a cell bound to a Zymosan particle while maintaining a healthy morphology throughout the imaging time-course. Imaging analysis was performed using FIJI software. The parameters for image processing were kept constant when comparing different data sets. The area of the phagosome was selected, cropped and opened in a new window. The threshold at different time points was measured by the analyze-measure function of FIJI. The values for the GFP-threshold of the phagosome area were divided by the values for the GFP-threshold of the whole cell at the corresponding time points.

### Dectin-1 and CD45 displacement assay

RAW 264.7 WT and Sptlc2^*-/-*^ cells were seeded on glass coverslips in 12-well plates and grown to 70-90% confluency overnight. Cells were transfected with pMXsIP Dectin-1-GFP using Lipofectamine 3000 (Invitrogen) according to the manufacturer’s instructions. After 24 hrs, cells were washed three times with ice-cold PBS, kept on ice for 5 min, and inoculated with Alexa Fluor 594-conjugated Zymosan A in chilled serum-free DMEM at a ratio of 10 beads per cell. After centrifugation for 5 min at 250 x g and 4°C, cells were incubated for 2 min at 37°C, washed three times with ice-cold PBS to remove unbound Zymosan, then fixed for 15 min with 4% paraformaldehyde. Images were collected using an LSM880 confocal microscope using a PLAN APO 63x oil-immersion objective and AIRY SCAN. Image analysis was performed using FIJI. Briefly, a region-of-interest was defined around the Zymosan particle, and Pearson’s correlation coefficient between Dectin-1-GFP signal and anti-CD45-APC signal was calculated using the colocalization function.

### Flow cytometry

RAW 264.7 wild type and Sptlc2^-/-^ cells were rinsed with PBS and lifted from the flask using treatment with trypsin. Fc receptor was blocked with CD16/32 antibody (1:100) and cells were stained with ZombieAqua Live/Dead (1:200). Cells were then stained with either CD45-APC (1:200), or Dectin-1-FITC (1:50) for 30 min. Finally, cells were fixed for 20 mins in 4% paraformaldehyde and analyzed using an LSR-II flow cytometer. Cells independently stained with ZombieAqua, CD45-APC, and Dectin-1-FITC were used for compensation. Analysis was performed using FlowJo software.

## Acknowledgements

This work was supported by the Medical Research Foundation and National Institutes of Health (3R21AI124225-01A1) to F.G.T., and the Deutsche Forschungsgemeinschaft (Projects SFB944-P14 and HO3539/1-1) to J.C.M.H. We would like to thank Dr. Hidde Ploegh (Boston Children’s Hospital, Boston, MA), Dr. Arturo Zychlinsky (Max Planck Institute of Biochemistry, Berlin, Germany), and Dr. David Johnson (Oregon Health and Science University, Portland, OR) for kindly providing reagents, cell lines, viruses, and plasmid constructs essential to the completion of this study. We would also like to thank the members of the OHSU Advanced Light Microscopy Core and the OHSU Flow Cytometry Shared Resource for assistance and training in data collection and analysis – in particular, we would like to thank Dr. Stefanie Kaech Petrie, Matthew Lewis, and Sara Christensen. We would finally like to thank Madeleine Faucher, Marie Foss, and Jossef Osborn who kindly and patiently assisted in the editing and preparation of this work.

## Author Contributions

P.N. and G.G. designed and performed the majority of experiments, and together prepared the manuscript. H.L. optimized and performed fluorescent microscopy colocalization and assisted in figure preparation. A.R., V.R., and F.P. created and characterized CRISPR knockout cell lines. J.C.M.H. provided intellectual contributions in experimental design and data interpretation. F.G.T. directed the research, interpreted the data, and edited the manuscript.

## Movie Legends

**Movie 1.** A movie showing live confocal microscopy imaging of wild type RAW 264.7 cells transfected with the Rac1/Cdc42 biosensor PAK(PBD)–GFP and infected Zymosan-conjugated with Alexa Flour 594 (shown in red) at a MOI of 10.

**Movie 2.** A movie showing live confocal microscopy imaging of Sptlc2^-/-^ RAW 264.7 cells transfected with the Rac1/Cdc42 biosensor PAK(PBD)–GFP and infected Zymosan-conjugated with Alexa Flour 594 (shown in red) at a MOI of 10.

**Movie 3.** A movie showing live confocal microscopy imaging of wild type RAW 264.7 cells transfected with the F-actin biosensor LifeAct-GFP and infected with Zymosan-conjugated with Alexa Flour 594 (shown in red) at a MOI of 10.

**Movie 4.** A movie showing live confocal microscopy imaging of Sptlc2^-/-^ RAW 264.7 cells transfected with the F-actin biosensor LifeAct-GFP and infected with Zymosan-conjugated with Alexa Flour 594 (shown in red) at a MOI of 10.

**Movie 5.** A movie showing live confocal microscopy imaging of wild type RAW 264.7 cells transfected with the PIP2 biosensor PH(PLCδ)-GFP and infected with Zymosan-conjugated with Alexa Flour 594 (shown in red) at a MOI of 10.

**Movie 6.** A movie showing live confocal microscopy imaging of Sptlc2^-/-^ RAW 264.7 cells transfected with the PIP2 biosensor PH(PLCδ)-GFP and infected with Zymosan-conjugated with Alexa Flour 594 (shown in red) at a MOI of 10.

**Movie 7.** A movie showing live confocal microscopy imaging of wild type RAW 264.7 cells transfected with the PIP3 biosensor PH(AKT)-GFP and infected with Zymosan-conjugated with Alexa Flour 594 (shown in red) at a MOI of 10.

**Movie 8.** A movie showing live confocal microscopy imaging of Sptlc2^-/-^ RAW 264.7 cells transfected with the PIP2 biosensor PH(AKT)-GFP and infected with Zymosan-conjugated with Alexa Flour 594 (shown in red) at a MOI of 10.

## Supplementary Results

**Supplemental Figure 1:**
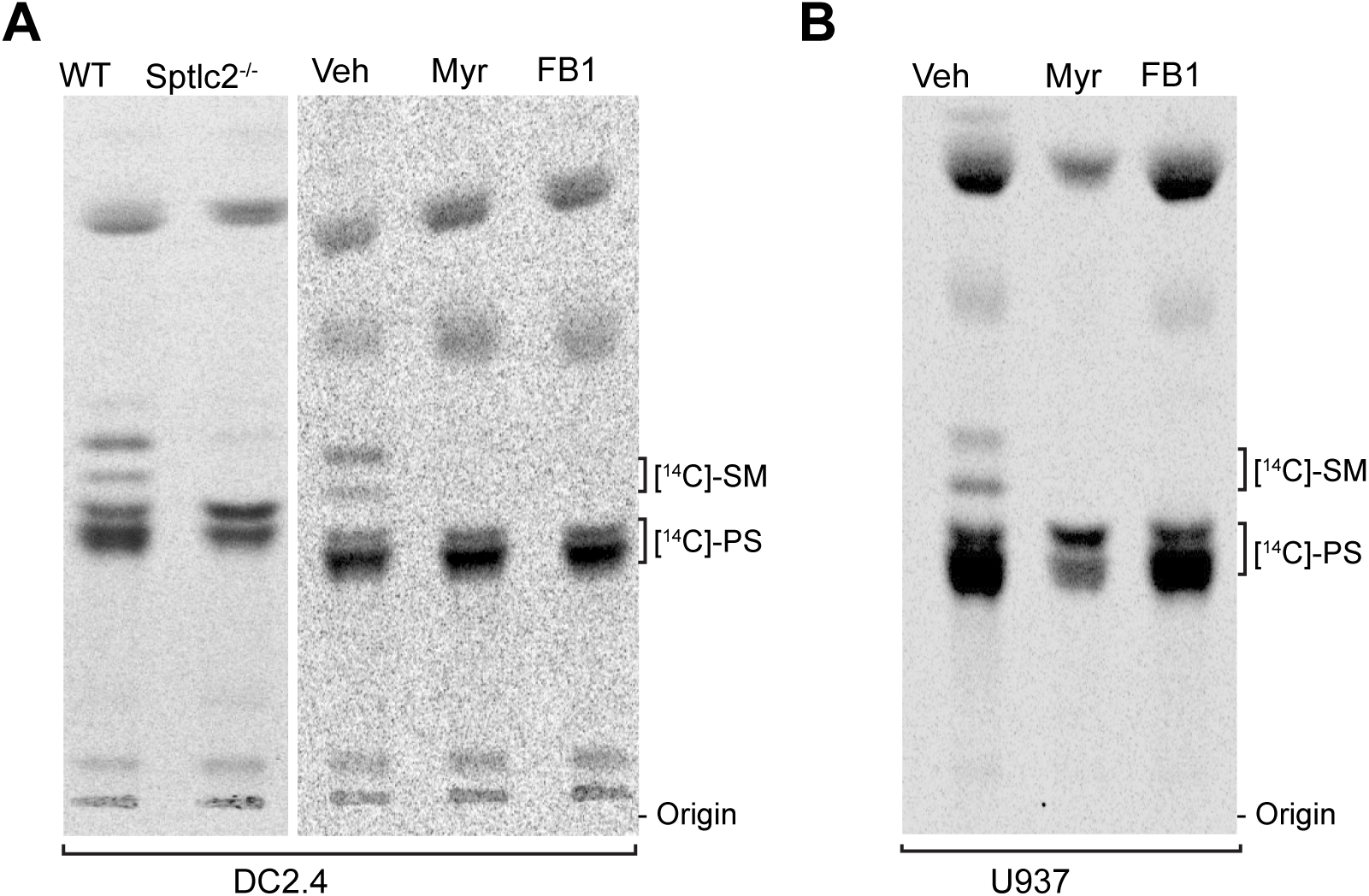
Genetic and small molecule inhibitors mediated depletion of sphingolipid in phagocytes. (A, B) Control, Sptlc2^-/-^ and inhibitor-treated cells were labeled with the sphingolipid precursor 3-L-[^14^C]-serine, and total lipids were extracted and analyzed by thin layer chromatography and autoradiography. All data are representative of 3 independent experiments.

**Supplemental Figure 2:**
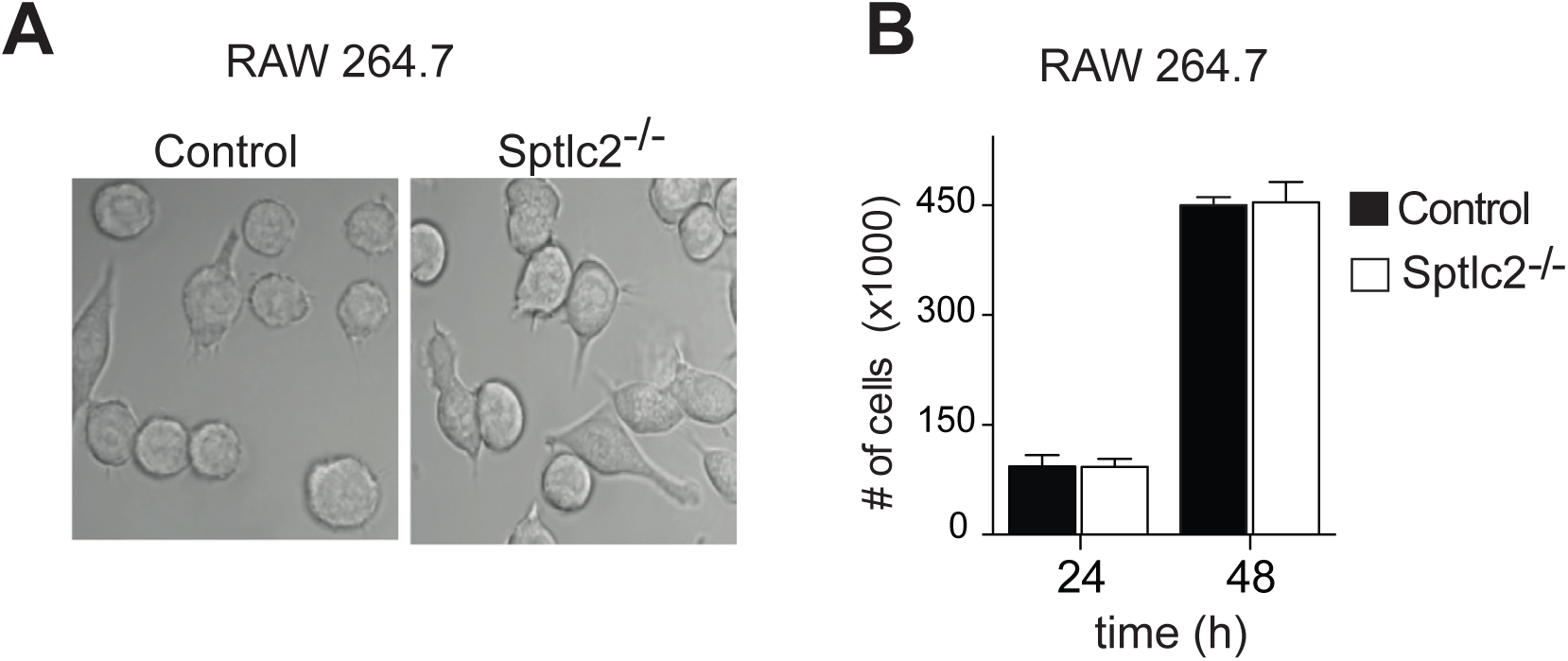
Depletion of sphingolipids has no significant effect on cell growth. (A) Microscopy images of control and Sptlc2^-/-^ RAW 264.7 macrophages, showing no morphological difference between the mutant and control cells. (B) The cell division of the control and Sptlc2^-/-^ RAW 264.7 macrophages at different time points was measured and presented.

**Supplemental Figure 3:**
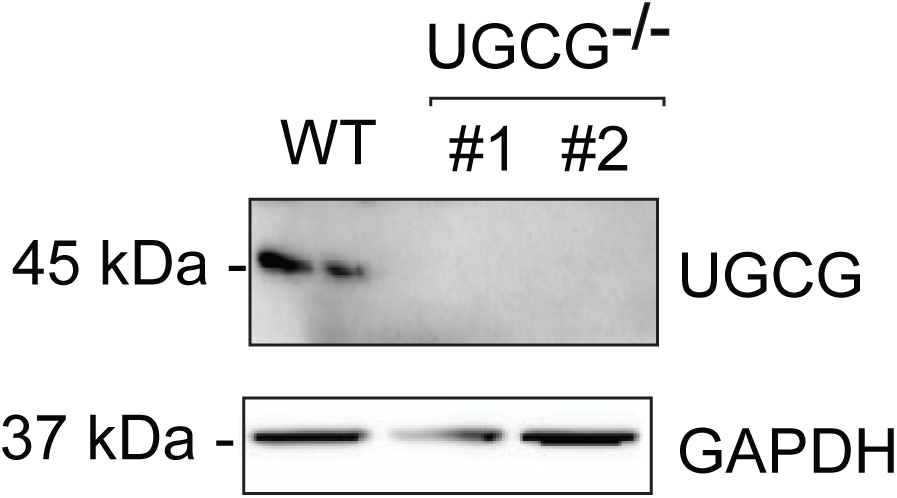
Western blot analysis of cell lysates collected from wildtype and two different CRISPR/Cas9 knockout THP-1 cell lines probed using anti-UGCG antibody (upper panel). Antibody against GAPDH was used as a loading control (lower panel).

